# CancerMIRNome: an interactive analysis and visualization database for miRNome profiles of human cancer

**DOI:** 10.1101/2020.10.04.325670

**Authors:** Ruidong Li, Han Qu, Shibo Wang, John M. Chater, Xuesong Wang, Yanru Cui, Lei Yu, Rui Zhou, Qiong Jia, Ryan Traband, Meiyue Wang, Weibo Xie, Dongbo Yuan, Jianguo Zhu, Wei-De Zhong, Zhenyu Jia

**Affiliations:** Department of Botany and Plant Sciences, University of California, Riverside, CA, USA; Graduate Program in Genetics, Genomics, and Bioinformatics, University of California, Riverside, CA, USA; College of Agronomy, Hebei Agricultural University, Baoding, China; Department of Anesthesiology, Perioperative and Pain Medicine, Stanford University School of Medicine, Stanford, CA, USA; Hubei Key Laboratory of Agricultural Bioinformatics, College of Informatics, Huazhong Agricultural University, Wuhan, China; Department of Urology, Guizhou Provincial People’s Hospital, Guizhou, China; Department of Urology, Guangdong Key Laboratory of Clinical Molecular Medicine and Diagnostics, Guangzhou First People’s Hospital, School of Medicine, South China University of Technology, Guangzhou, China; Urology Key Laboratory of Guangdong Province, The First Affiliated Hospital of Guangzhou Medical University, Guangzhou Medical University, Guangzhou, China; Macau Institute for Applied Research in Medicine and Health, Macau University of Science and Technology, Macau, China

## Abstract

MicroRNAs (miRNAs), which play critical roles in gene regulatory networks, have emerged as promising diagnostic and prognostic biomarkers for human cancer. In particular, circulating miRNAs that are secreted into circulation exist in remarkably stable forms, and have enormous potential to be leveraged as non-invasive biomarkers for early cancer detection. Novel and user-friendly tools are desperately needed to facilitate data mining of the vast amount of miRNA expression data from The Cancer Genome Atlas (TCGA) and large-scale circulating miRNA profiling studies. To fill this void, we developed CancerMIRNome, a comprehensive database for the interactive analysis and visualization of miRNA expression profiles based on 10,998 samples from 33 TCGA projects and 21,993 samples from 40 public circulating miRNome datasets. A series of cutting-edge bioinformatics tools and machine learning algorithms have been packaged in CancerMIRNome, allowing for the pan-cancer analysis of a miRNA of interest across multiple cancer types and the comprehensive analysis of miRNome profiles to identify dysregulated miRNAs and develop diagnostic or prognostic signatures. The data analysis and visualization modules will greatly facilitate the exploit of the valuable resources and promote translational application of miRNA biomarkers in cancer. The CancerMIRNome database is publicly available at http://bioinfo.jialab-ucr.org/CancerMIRNome.

## INTRODUCTION

miRNAs are a class of small endogenous non-coding RNAs of ∼22nt in length that negatively regulate the expression of their target protein-coding genes (1). It has been reported that miRNAs are involved in many biological processes, such as cell proliferation, differentiation, and apoptosis (2–5). Mounting evidence has demonstrated that miRNAs are dysregulated in various types of human cancer (6–8), which may be leveraged as expression signatures for cancer diagnosis and prognosis. Circulating miRNAs represent the miRNAs that are secreted into extracellular body fluids, where they are incorporated into extracellular vesicles (EVs), such as shed microvesicles (sMVs) and exosomes, or in apoptotic bodies, or form complexes with RNA binding proteins, such as Argonates (AGOs). These protected circulating miRNAs remain in remarkably stable forms, rendering potential cancer biomarkers for non-invasive early detection or tissue-of-origin localization (9–11).

The vast amount of miRNA expression data in TCGA as well as data from many large-scale circulating miRNA profiling studies are readily available for the discovery and validation of miRNA biomarkers for cancer diagnosis and prognosis (12–14). Two online resources – OncomiR (15) and OMCD (16) have been developed for the exploring of miRNA expression profiles to identify dysregulated miRNAs associated with clinical characteristics of cancer based on TCGA data. While the functional modules provided by OncomiR and OMCD are very useful, certain important functions are lacking for the comprehensive analysis of cancer miRNome data, such as functional enrichment analysis of miRNA targets, identification of diagnostic biomarkers, development of machine learning-based prognostic models, dimensionality reduction analysis, etc. In addition, the functionalities provided in the existing databases are relatively simple. For example, although differential expression (DE) analysis is supported by both databases, users cannot define their own groups for comparison, e.g., late tumor stages (III + IV) vs. early tumor stages (I + II). Note that DE analysis is the only analytical function available in OMCD. Univariate Cox Proportional-Hazards (CoxPH) survival analysis can be performed in OncomiR, but the commonly applied statistics in survival analysis - hazard ratio (HR) and confidence interval (CI) - are not reported to the end users. Moreover, data visualization and export are not well supported by OncomiR or OMCD, which constrains their broad application. Most importantly, only miRNA expression profiling data of tumor or tumor-adjacent normal tissues from TCGA were included in the databases. Sophisticated and user-friendly web tools are desperately needed to not only facilitate the exploit of TCGA miRNome data, but also allow for data mining of the valuable circulating miRNome data resources to promote translational research on cancer miRNAs. To fill this void, we developed CancerMIRNome, an integrated database for the interactive analysis and visualization of miRNA expression profiling data of human tissues and body fluids of cancer patients with 10,998 samples from 33 TCGA projects and 21,993 samples of 32 cancer types from 40 circulating miRNA profiling studies (Figure 1).

**Figure 1.**
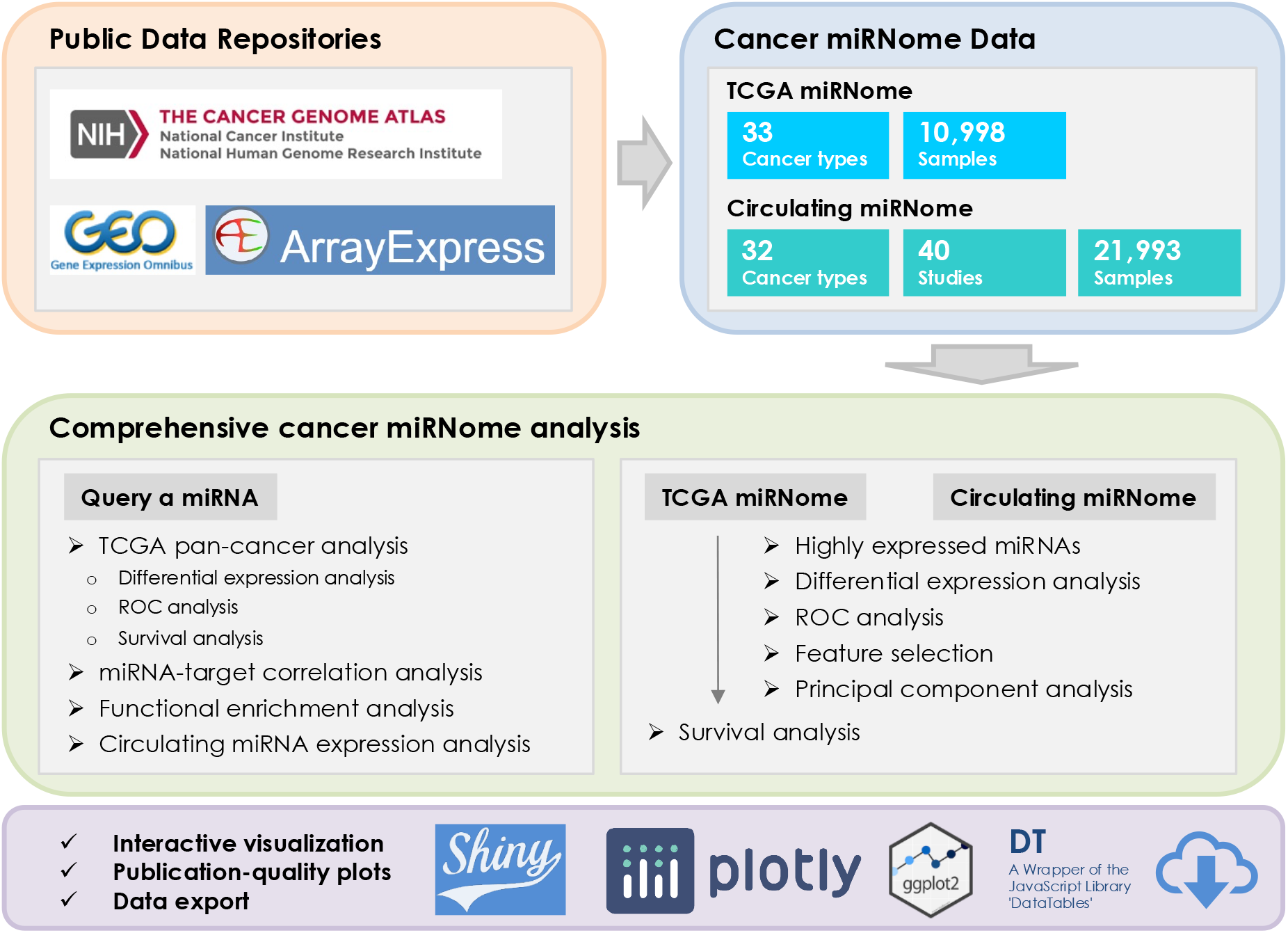
Overview of the CancerMIRNome database.

CancerMIRNome is the most comprehensive database of miRNome profiles of human cancer to date and it provides a suite of advanced functions for (i) pan-cancer characterization of a miRNA of interest across multiple cancer types, including differential expression analysis, survival analysis, miRNA-target correlation analysis, functional enrichment analysis, and circulating miRNA expression analysis; and (ii) comprehensive miRNome data analysis, including the identification of highly expressed miRNAs, selection of diagnostic miRNA biomarkers based on differential expression analysis, receiver operating characteristic (ROC) curve, and least absolute shrinkage and selection operator (Lasso) algorithm, dimensionality reduction analysis, univariate survival analysis, and the development of prognostic models. Advanced visualizations are supported to produce publication-quality vector images in PDF format. All processed data deposited in CancerMIRNome, including the normalized miRNA expression data and harmonized sample metadata for each dataset can be easily downloaded, allowing for further analysis by the end users (Figure 1).

## DATA COLLECTION AND PROCESSING

### TCGA miRNome profiles

A unified bioinformatics pipeline based on the R/Bioconductor package GDCRNATools (19) was developed to download and process the mature miRNA expression data and clinical data of 33 cancer types in TCGA. Isoform expression quantification data of miRNA-Seq were downloaded from National Cancer Institute (NCI) Genomic Data Commons (GDC) using the *gdcRNADownload* function. The expression data from the same project were merged to a single expression matrix using the *gdcRNAMerge* function in the R package GDCRNATools, followed by a normalization with the Trimmed Mean of M-values (TMM) normalization method implemented in the R package edgeR (20). Clinical information including age, tumor stages, overall survival, etc. were retrieved from the XML file of each sample using the *gdcClinicalMerge* function in GDCRNATools.

### Circulating miRNome profiles

An extensive search for circulating miRNA expression profiles of human cancer was performed in public data repositories, including the National Center for Biotechnology Information (NCBI) Gene Expression Omnibus (GEO) and ArrayExpress. A total of 40 public circulating miRNA expression datasets with over 1000 miRNAs profiled in each dataset were identified by searching the keywords ‘circulating’, ‘blood’, ‘serum’, ‘plasma’, ‘extracellular vesicle’, or ‘exosome’, in combination with ‘miRNA’ or ‘microRNA’, and with ‘cancer’, ‘tumor’, or ‘carcinoma’. Both the normalized miRNA expression data and sample metadata from GEO were downloaded by the *getGEO* function in the R package GEOquery (21). The processed data in ArrayExpress were downloaded directly from the website. miRNA annotation information from miRBase release 10.0 to the latest release 22.1 were integrated and the miRNA names in each version were mapped to the stable miRNA accession numbers (beginning with MIMAT) for the harmonization of miRNA identifiers in all the circulating miRNome datasets. If multiple probes matched to the same miRNA accession number, only the one with the maximum interquartile range (IQR) for the miRNA expression values was kept. The log2 transformation may be performed on the miRNA expression data if it hasn’t been applied to the original data. Metadata of the samples were harmonized using a custom script (available at https://github.com/rli012/CancerMIRNome) followed by a careful manual curation.

## DATABASE CONTENT AND USAGE

A series of cutting-edge bioinformatics tools and functions have been packaged in CancerMIRNome, allowing for the pan-cancer analysis of a miRNA of interest across multiple cancer types and the comprehensive analysis of cancer miRNome profiles. Many advanced visualization methods are supported to facilitate the interpretation of the results. Moreover, all processed data deposited in CancerMIRNome, the outputs from the data analyses including tables and high-resolution figures, as well as the data that are used to generate the figures can be easily downloaded from the CancerMIRNome database. The web interface of CacnerMIRNome is highly intuitive to exploit and a step-by-step tutorial is provided for users.

### Query a miRNA of interest

Users can query a miRNA of interest by typing the miRNA accession number, miRNA ID of the latest miRBase release 22.1 (22), or previous miRNA IDs in the ‘Search a miRNA’ field and selecting this miRNA from the dropdown list. In addition to the general information such as IDs and sequence of the queried miRNA, external links to five miRNA-target databases, including ENCORI (23), miRDB (24), miTarBase (23), TargetScan (26), and Diana-TarBase (27), are also provided to facilitate the exploring of the miRNA using different online resources.

The single miRNA analysis modules include: (i) pan-cancer differential expression (DE) analysis, receiver operating characteristic (ROC) analysis, and Kaplan Meier (KM) survival analysis in TCGA; (ii) DE, ROC, an KM survival analysis in an individual TCGA project; (iii) miRNA-target correlation analysis; (iv) functional enrichment analysis of the miRNA targets; and (v) circulating miRNA expression analysis (Figure 2).

**Figure 2.**
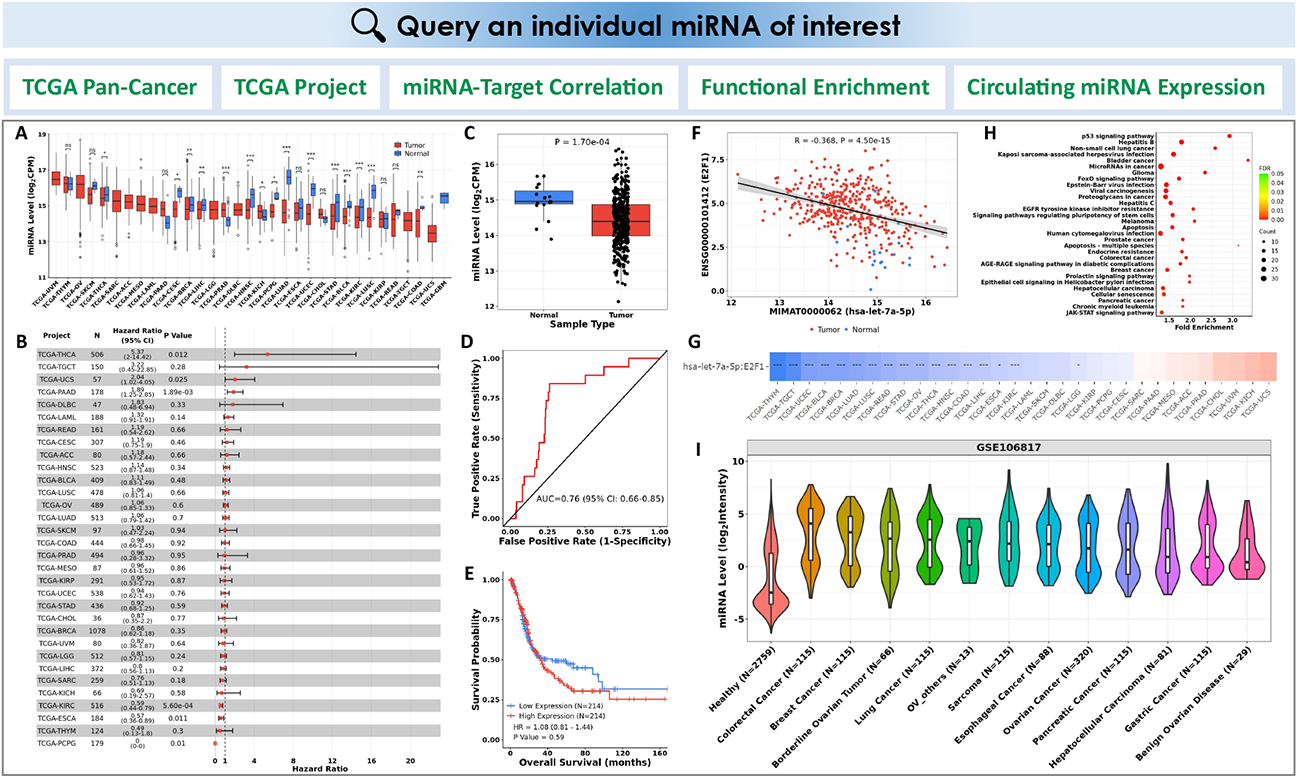
CancerMIRNome outputs from the query of a miRNA of interest. (**A**) Pan-cancer differential expression analysis across all TCGA projects. (**B**) A forest plot visualizing pan-cancer survival analysis across all TCGA projects. (**C**) Boxplot of the miRNA expression in tumor and normal samples from the selected TCGA project. (**D**) An ROC curve illustrating the diagnostic ability of the miRNA in the selected TCGA project. (**E**) Kaplan Meier analysis of overall survival between tumor samples with high and low expression of the miRNA of interest defined by its median expression value in the selected TCGA project. (**F**) Correlation analysis of the miRNA with one of its targets in a TCGA project. (**G**) An interactive heatmap visualizing the miRNA-target correlations across all TCGA projects. (**H**) A bubble plot visualizing the functional enrichment of target genes for the miRNA of interest. (**I**) A violin plot visualizing the circulating miRNA expression in a selected circulating miRNome dataset of human cancer.

#### 1. Pan-cancer analysis

Pan-cancer analysis can be performed on TCGA data across 33 cancer types to investigate the dysregulation of a miRNA of interest and its potential to be used as diagnostic and prognostic markers in cancer. For example, differential expression (DE) analysis is conducted to determine if the miRNA of interest is differentially expressed between tumor and normal samples, the area under the ROC curve (AUC) is computed to measure the performance of the miRNA biomarker in differentiating tumor samples from normal samples, while the KM analysis of overall survival (OS) in patients with high *versus* low expression levels of the miRNA in the corresponding primary tumor samples can be performed to assess its prognostic ability. The expression levels and the statistical significance of the miRNA in all the TCGA projects can be visualized in a box plot (Figure 2A). A forest plot displaying the number of tumor and normal samples, AUC, and 95% confidence interval (CI) of the AUC for each cancer type in TCGA is used to visualize the result of pan-cancer ROC analysis. The forest plot is also generated to visualize the pan-cancer KM survival analysis by showing the number of tumor samples, hazard ratio (HR), 95% confidence interval (CI) of the HR, and p value for each TCGA project (Figure 2B).

#### 2. miRNA analysis in an individual TCGA project

Th DE, ROC, and KM survival analyses can also be implemented for a selected TCGA project to show more detailed information about the dysregulation of the miRNA and its associated diagnostic/prognostic power for a specific cancer type. A box plot with miRNA expression between tumor and normal samples, an ROC curve, and a KM survival curve for the selected project will be displayed (Figure 2 C-E).

#### 3. miRNA-target correlation analysis

The vast amount of miRNA and mRNA expression data from over 10,000 samples in TCGA provides a tremendous opportunity to systematically investigate the miRNA-mRNA associations in cancer. Pearson correlation between a miRNA and its target genes can be evaluated in CancerMIRNome to uncover the relationship of their expression intensities in the TCGA datasets. The miRNA-target interactions are based on miRTarBase 2020 – an experimentally validated miRNA-target interactions database. The expression correlations between a miRNA and all of its targets in a selected TCGA project are listed in an interactive data table. Users can select an interested interaction between miRNA and mRNA target in the data table to visualize a scatter plot showing their expression pattern and correlation metrics (Figure 2F). An interactive heatmap is also available to visualize and compare the strength of miRNA-target correlations across all the 33 cancer types in TCGA (Figure 2G).

#### 4. Functional enrichment analysis of miRNA targets

The identification of biological pathways in which the miRNAs are involved is critical to understand the regulatory roles of miRNAs in human cancer. In CancerMIRNome, functional enrichment analysis of the target genes for a miRNA of interest can be conducted using clusterProfiler (28) with support of many pathway/ontology knowledgebases, including Kyoto Encyclopedia of Genes and Genomes (KEGG) (29), Gene Ontology (GO) (30), Reactome (31), Disease Ontology (DO) (32), Network of Cancer Gene (NCG) (33), DisGeNET (34), and Molecular Signatures Database (MSigDB) (35). A data table will be created to summarize the significantly enriched pathways/ontologies with the number and proportion of enriched genes, the significance levels, as well as the gene symbols in the pathway/ontology terms. The top 30 pathways/ontologies can be visualized as bar plot and bubble plot (Figure 2H).

#### 5. Circulating miRNA expression

Expression of an interested miRNA in whole blood, serum, plasma, EVs, or exosomes from both healthy and cancer patients can be conveniently explored in the 40 circulating miRNome datasets. Users can select a dataset from the interactive data table for an analysis of the circulating miRNA expression, through which a violin plot is displayed for visualization and comparison (Figure 2I).

### Comprehensive miRNome analysis

CancerMIRNome is equipped with well-designed functions for the comprehensive miRNome profiling data analysis in each of the 33 TCGA projects and the 40 circulating miRNome datasets. In spite of some slight differences, the bioinformatics techniques for the analysis and visualization of TCGA miRNome data and circulating miRNome data are largely the same, therefore, we describe the functions in both tabs, i.e., ‘TCGA miRNome’ and ‘Circulating miRNome’, at the same time in this section.

When a dataset is selected, the summary of important clinical features of patients, including sample type, tumor stages, ages, overall survival, etc., will be displayed for this TCGA project (Figure 3 A-C), whereas, the platform, the statistics of the cancer types, and an embedded webpage (from either GEO or ArrayExpress) housing the public dataset will be shown for the circulating miRNome dataset.

**Figure 3.**
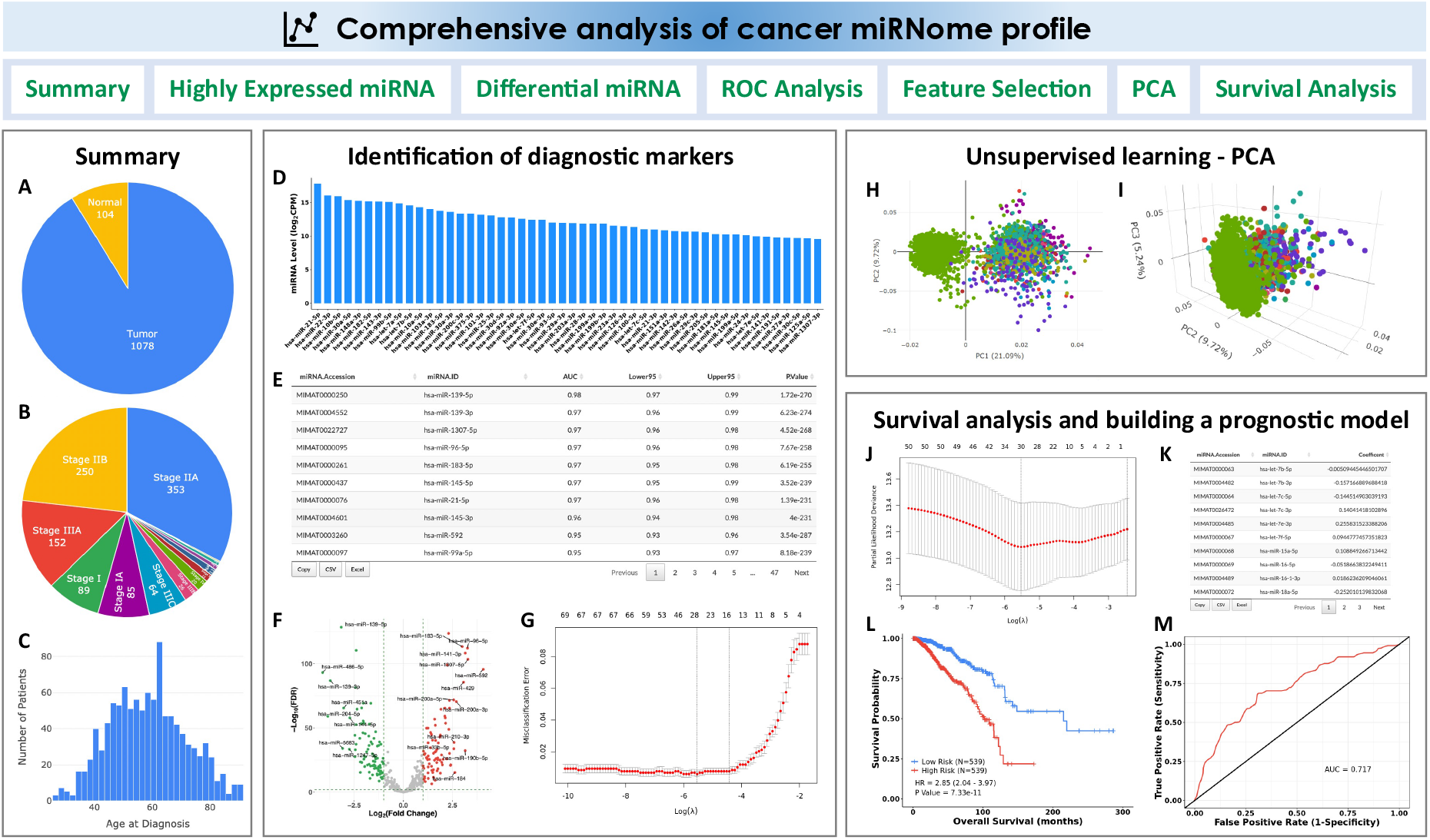
CancerMIRNome outputs from the comprehensive analysis of a miRNome dataset. (**A**) Pie plot showing the statistics of sample type for a TCGA project. (**B**) Pie plot visualizing the statistics of clinical stage for a TCGA project. (**C**) Distribution of age at diagnosis for the patients in a TCGA project. (**D**) Bar plot of top 50 highly expressed miRNAs. (**E**) A data table for the diagnostic markers identified by ROC analysis. (**F**) A volcano plot visualizing the differentially expressed miRNAs between two user-defined groups. (**G**) Selection of the most-relevant diagnostic miRNA biomarkers using Lasso. (**H**) 2D interactive visualization of principal component analysis result using the first two principal components. (**I**) 3D interactive visualization of principal component analysis result using the first three principal components. (**J**) Selection of prognostic miRNA biomarkers using the Cox-Lasso technique to develop a prognostic model. (**K**) Coefficients of the selected miRNAs in the prognostic model. (**L**) Kaplan Meier survival analysis evaluating the prognostic ability of the miRNA expression-based prognostic model. (**M**) Time-dependent ROC analysis evaluating the prognostic ability of the model.

The comprehensive miRNome data analysis modules include: (i) identification of highly expressed miRNAs; (ii) DE analysis between two user-defined groups; (iii) ROC analysis to evaluate the classification performance of the highly expressed miRNAs; (iv)identification of diagnostic miRNA markers based on a machine learning algorithm; (v) principal component analysis for dimensionality reduction; and (vi) survival analysis and miRNA expression-based prognostic model development (only for TCGA projects where the overall survival data are available) (Figure 3).

#### 1. Highly expressed miRNAs

miRNAs with relatively high abundances may be more reliable, robust and practical to be used as diagnostic or prognostic biomarkers in clinical settings. The highly expressed miRNAs are identified if their counts per million (CPM) are greater than 1 in more than 50% of the samples in a TCGA project or if their abundances are ranked among the top 500 miRNAs in a circulating miRNome dataset. The top 50 highly expressed miRNAs are visualized in a bar plot based on their median expression values (Figure 3D).

#### 2. ROC analysis

The ROC analysis can be carried out to screen the highly expressed miRNAs for the identification of diagnostic biomarkers to distinguish tumor samples from normal samples in a TCGA dataset or distinguish liquid biopsy samples from cancer patients and healthy donors in a circulating miRNome dataset. All the miRNAs are ranked in an output data table based on their AUC values (Figure 3E).

#### 3. DE analysis

The DE analysis of miRNAs for a TCGA project allows users to identify dysregulated miRNAs that are associated with tumor initiation or progression by comparing the case and control groups. Circulating miRNAs with elevated expression levels can also be identified by DE analysis, which may be used as diagnostic biomarkers for non-invasive cancer detection. The highly expressed miRNAs in a dataset can be compared between two user-defined groups for identifying the significant DE miRNAs (Figure 3F). For TCGA projects, clinical variables, such as sample type, tumor stages, etc., may be utilized to group samples for comparison. For example, the DE analysis can be performed not only between tumor and normal samples, but also between patients at early and late tumor stages. The samples in the circulating miRNome datasets are mainly grouped by disease status or cancer types for DE analysis, e.g., lung cancer versus healthy, or lung cancer versus non-cancerous lung disease, etc. The R package limma (36) is implemented for the DE analysis.

#### 4. Machine learning-based feature selection

CancerMIRNome provides a machine learning algorithm - least absolute shrinkage and selection operator (Lasso) (37, 38) to detect miRNAs with diagnostic power and develop a classification model based on the expression values of the miRNA signature for cancer diagnosis. The Lasso regression uses the L1 regularization technique to shrink coefficients of the insignificant features to zero. The tuning parameter lambda (*λ*), which controls the overall strength of the L1 penalty, is determined based on a built-in 10-fold cross-validation. When a dataset is selected, the machine learning models are trained using the expression data of the highly expressed miRNAs to classify different types of samples, e.g., tumor versus normal samples for a TCGA project or serum samples from cancer patients versus healthy donors in a selected circulating miRNome dataset. The cross-validation curve is plotted (Figure 3G) and the coefficients for the most relevant features (miRNAs) at the selected value of *λ* that gives minimum mean cross-validated error are provided in a data table.

#### 5. Principal component analysis

The commonly employed unsupervised learning algorithm, principal component analysis (PCA), can be utilized for dimensionality reduction to analyze the high-dimensional miRNome expression profiling data for any selected dataset such that all patient samples may be visualized in a 2D or 3D interactive plot using the first two or three principal components, respectively (Figure 3H and Figure 3I).

#### 6. Survival analysis

Three survival analysis modules were developed in CancerMIRNome for the identification of prognostic miRNA biomarkers and development of miRNA expression-based prognostic models, including (i) univariate CoxPH regression analysis and KM survival analysis, (ii) creation of pre-built prognostic models using the regularized Cox regression model with Lasso penalty (Cox-Lasso) algorithm (39), and (iii) development of prognostic models based on the user-provided miRNA signatures. The data tables for the univariate CoxPH and KM survival analyses of all the highly expressed miRNAs in the selected TCGA project are provided. The pre-built prognostic model for each cancer type in TCGA was developed by jointly analyzing the miRNAs that are significant in the univariate CoxPH analysis. The coefficients of the most relevant miRNAs are provided, whereas those of the irrelevant variables were shrink to zero by the Cox-Lasso algorithm (Figure 3J and 3K). The prognostic model, which is a linear combination of the finally selected miRNA variables with the Cox-Lasso-derived regression coefficients, will be used to calculate a risk score for each patient. All the patients will be dichotomized into either a high-risk group or a low-risk group based on the median risk value for the cohort. The KM survival analysis and time-dependent ROC analysis can be performed to evaluate the prognostic ability of the miRNA-based prognostic model (Figure 3L and Figure 3M). Besides the pre-built models, CancerMIRNome also provides a module allowing for users to submit their own miRNA expression signatures of interest to build prognostic models using three survival analysis methods, including multivariate CoxPH, Cox-Lasso, and Cox regression model regularized with ridge penalty (Cox-Ridge) (39, 40).

### Data download

All the processed data, including the 33 TCGA miRNome datasets, the 40 circulating miRNome datasets of human cancers, and the integrated miRNA annotation data can be downloaded easily on the ‘Download’ page of CancerMIRNome. The ExpressionSet class is used for the miRNA expression data and metadata of the miRNome datasets. The miRNA annotation data includes the miRNA accession number, miRNA name, and miRNA sequence from the latest miRBase release 22.1, and the previous miRNA names from miRBase release 10.0 to release 21. The data are downloaded as RDS files, which can be easily imported into R. Moreover, the outputs from the data analyses including tables, high-resolution figures, and the data that are used to generate the figures are all exportable.

## SUMMARY AND FUTURE DIRECTIONS

In this study, we present to the cancer research community a user-friendly web tool, CancerMIRNome, for the interactive analysis and visualization of miRNome profiles of human cancer by leveraging 10,998 tumor and normal samples from 33 TCGA projects and 21,993 samples of 32 cancer types from 40 public circulating miRNA profiling studies. A suite of well-designed functions is provided to facilitate data mining at both the miRNA level and the miRNome level (or dataset level). For example, a comprehensive characterization of a miRNA of interest, including pan-cancer DE analysis, ROC analysis, survival analysis, miRNA-target correlation analysis, functional enrichment analysis, and circulating miRNA expression analysis, may be simply carried out by querying this miRNA on the CancerMIRNome webpage. This is tremendously helpful when users are interested in the expression and function of a miRNA across multiple cancer types; otherwise, one has to download, process and analyze the expression data and clinical data from all the TCGA projects to reach the same results, which requires high-level bioinformatics programming skills and substantial time and effort. Advanced visualizations are supported in CancerMIRNome and the publication-quality vector images can be easily created and downloaded. Moreover, all of the data and results are exportable, allowing for further local analyses by the end users. While CancerMIRNome is diligently serving the cancer research community, we are open to any feedback from users and will constantly maintain and improve this database. New datasets, analytical methods, and visualization functions will be included in CancerMIRNome as soon as they become available. We expect that CancerMIRNome would become a valuable online resource for a comprehensive analysis of cancer miRNome data not only for experimental biologists, but also for bioinformatics scientists in the field.

## AVAILABILITY

The CancerMIRNome database is publicly available at http://bioinfo.jialab-ucr.org/CancerMIRNome. The source code for processing miRNome data and building the database is available at https://github.com/rli012/CancerMIRNome. All the processed data deposited in CancerMIRNome can be downloaded easily on the ‘Download’ page of the database.

## FUNDING

This work was supported by Z.J.’s UC Riverside Faculty Start-up Fund, UC Cancer Research Coordinating Committee Competition Award, UC Academic Senate CoR Research Grant and United States Department of Agriculture (2019-67022-29930). D.Y. and J.Z. were supported by the National Natural Science Foundation of China (81660426), Science and Technology Project of Guizhou Province in 2017 ([2017]5803), the High-level innovative talent project of Guizhou Province in 2018 ([2018]5639), and the Science and Technology Plan Project of Guiyang in 2019 ([2019]2-15). R. Z. and W.Z. were supported by the grants from National Natural Science Foundation of China (82072813, 8157142), Guangzhou Municipal Science and Technology Project (201803040001).

## CONFLICT OF INTEREST

The authors declare that they have no competing interests.

## Notes

### Competing Interest Statement

The authors have declared no competing interest.

